# Rad9/53BP1 promotes crossover recombination DNA repair by limiting the Sgs1 and Mph1 helicases

**DOI:** 10.1101/638791

**Authors:** Matteo Ferrari, Chetan C. Rawal, Samuele Lodovichi, Achille Pellicioli

**Affiliations:** Dipartimento di Bioscienze, Università degli Studi di Milano, Via Celoria 26, 20131 Milano, Italia; Developmental Biology Program, Memorial Sloan Kettering Cancer Center, New York, NY 10065 USA; Department of Molecular and Computational Biology, University of Southern California, Los Angeles, CA 90089 USA

## Abstract

A DNA double strand break (DSB) is primed for homologous recombination (HR) repair through the nucleolytic processing (resection) of its ends, leading to the formation of a 3′ single-stranded DNA (ssDNA). Generation of the ssDNA is accompanied by the loading of several repair factors, including the ssDNA binding factor RPA and the recombinase Rad51. Then, depending upon the availability and location of a homologous sequence, different types of HR mechanisms can occur. Inefficient or slow HR repair results in the activation of the DNA damage checkpoint (DDC)^1^. In budding yeast, the 53BP1 ortholog Rad9 acts as a scaffold, mediating signal from upstream kinases Mec1 and Tel1 (ATR and ATM in human) to downstream effectors kinases Rad53 and Chk1 (CHK2 and CHK1 in human). In addition to its role in DDC, Rad9 limits DSB resection ^2^. Remarkably, this function is conserved in 53BP1, also being implicated in cancer biology in human cells ^3,4^.

Here we show that Rad9 limits the recruitment of the helicases Sgs1 and Mph1 on to a DSB, promoting Rad51-dependent recombination with long track DNA conversions, crossovers and break-induced replication (BIR). This regulation couples the DDC with the choice and effectiveness of HR sub-pathways, and might be critical to limit genome instability with implication for cancer research.

## Main

Rad9 is the 53BP1 ortholog in *S. cerevisiae* (budding yeast). It plays fundamental role in DNA damage signalling, mediating phosphorylation from upstream kinases Mec1/ATR and Tel1/ATM. In addition to this function, we described a DDC-independent role of Rad9 in the regulation of DSB resection, limiting repair through single strand annealing (SSA). Indeed, the Rad9 binding to a DSB reduces the Exo1 and Dna2-Sgs1 recruitment, affecting the long-range resection^2,5,6^. However, it is not understood how the formation of the 3′-end filament is coupled with the selection of appropriate HR sub-pathway, and whether Rad9 might have a role to determine that choice.

To investigate further the role of Rad9 in DSB repair, we performed a genetic assay with a diploid system, which allows to study the repair of DSB through gene conversion (GC) and break-induced replication (BIR), after the induction of a DSB by I-*Sce*I in chromosome XV^7^. Importantly, this system not only measures frequencies of crossover (CO) and noncrossover (NCO) in DSB repair, but also distinguishes between short and long tract gene conversion events (Fig. 1a). Upon I-*Sce*I induction, in the absence of *RAD9* we observed a striking increase in the percentage of short track GC (white colonies in the assay) (Fig. 1b). By screening colonies for specific markers, we also found that *rad9*Δ cells have reduced frequency of both CO and BIR events (Fig. 1c, and Extended data Table 1). Based on these results, we hypothesized that Rad9 might control strand invasion-mediated mechanisms to repair a DSB, in addition to its function in limiting SSA.

**Fig. 1:**
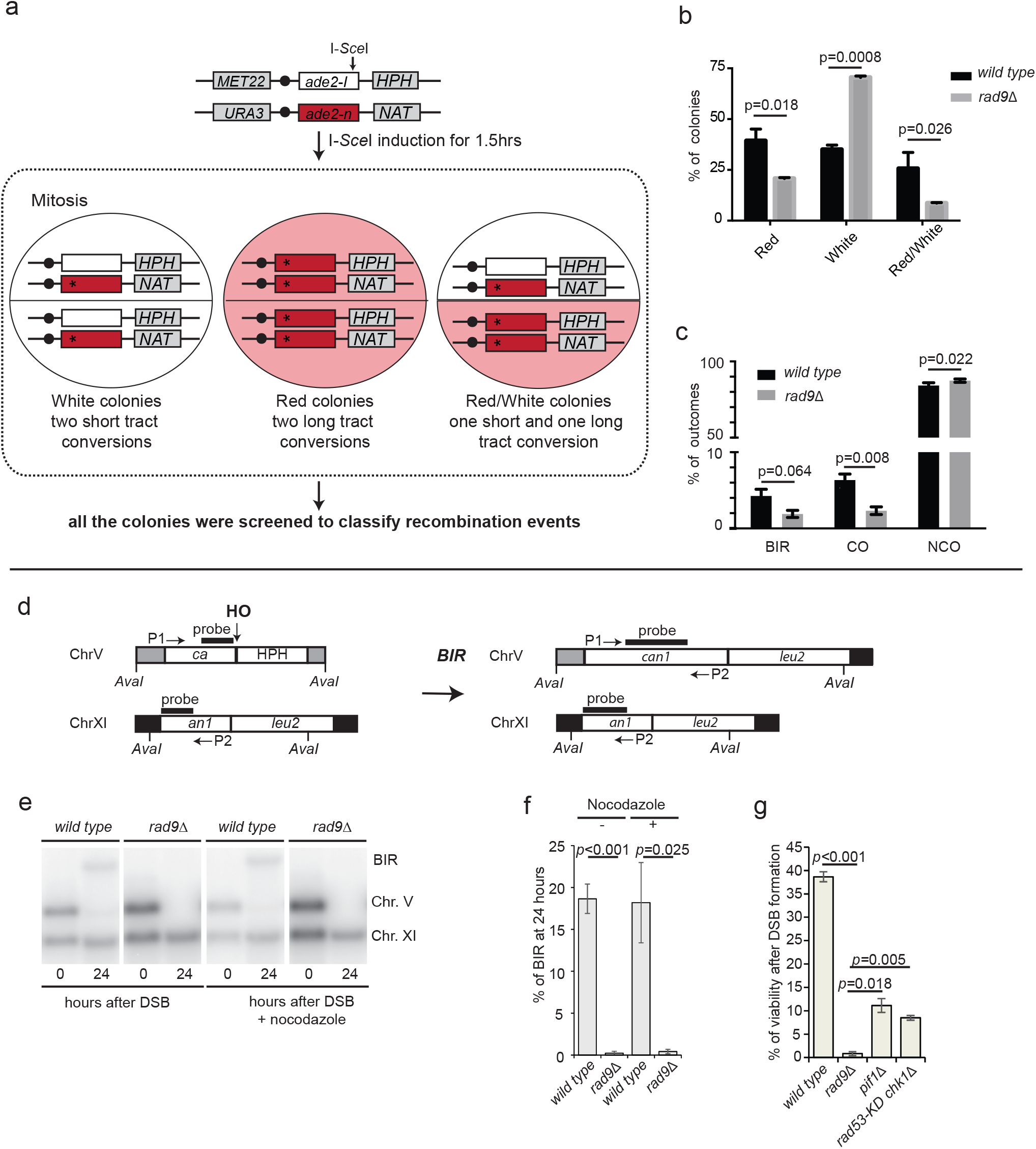
Rad9 promotes crossover recombination DSB repair. a, scheme of the genetic system to test recombination events in LSY2205-11C/LSY2543 background. b, Distribution of the recombinant colony types in the indicated LSY2205-11C and LSY2543 derivative diploid strains, with SEM of three experiments. Red (two long track conversions), White (two short track conversions) and Red/White (one long and one short track conversion) were determined from recombinant colonies after I-*Sce*I induction. c, Distribution of CO, NCO and BIR among all the colony types of the experiments in (b), with SEM of two independent diploids. d, scheme of the genetic system to test BIR in YRL92 background. e, Southern blot of *Ava*I-digested DNA to monitor DSB repair through BIR in the indicated YRL92 strains. f, Densitometric analysis of the BIR band of three experiments as in (e), with SEM. g, BIR efficiency measured by cell viability in the indicated YRL92 strains, with SEM of three experiments.

To study more precisely the role of Rad9 in the BIR process, we took advantage of an haploid genetic system engineered to test only this HR subpathway^8^. Briefly, one DSB is induced in chromosome V by the endonuclease HO and it is repaired by BIR thanks to a donor sequence on chromosome XI (Fig. 1d). Interestingly, analysing the DSB repair by Southern blotting, in exponentially growing or nocodazole-arrested cells (Fig. 1e, f), we found that *RAD9* deletion severely affected BIR, regardless of the cell cycle stage. Considering that DDC signalling to Pif1 helicase contributes to BIR^9^, we speculated that this would be the reason of the *rad9*Δ cells defect in the assay. However, after plating the cells in galactose to induce HO, we surprisingly observed higher cell lethality in *rad9*Δ cells with respect to *rad53*-K227A *chk1*Δ cells, that are defective in DDC signalling downstream from Rad9, and *pif1*Δ cells (Fig. 1g). Therefore, cells lacking *RAD9* would be defective in BIR for a reason other than deficient signalling to Pif1.

Then, we tested critical steps in Rad51-dependent HR in *rad9*Δ cells. First, by chromatin immunoprecipitation (ChIP), we found that the recombination factors Rpa1, Rad52 and Rad51 were efficiently recruited at the cut site in *rad9*Δ cells (Fig. 2a, b, c, and Extended data Fig. 1). However, their binding near to the break ends was significantly higher in the absence of *RAD9*, especially at later time points. Since the amount of ssDNA close to the break site in wild type and *rad9*Δ cells was similar (Extended data Fig. 2), the increased recruitment of Rpa1, Rad52 and Rad51 on the DSB was unlikely due to higher amount of available substrate in *rad9*Δ cells. Instead, we hypothesized that the loading and oligomerization of Rad9 protein on the DSB might physically dampen the recruitment of the recombination factors, similarly to its role in limiting nucleases for DSB resection^2,5,6^. To test this hypothesis, we expressed from a plasmid the wild type Rad9 or the two protein variants Rad9-2Ala and Rad9-7xA, both reducing the Rad9 binding and oligomerization on the DSB^5,10^, in *rad9*Δ cells. We found that both the Rad9 variants, contrary to the wild type form, did not completely rescue the lethality of *rad9*Δ cells in the BIR assay (Extended data Fig. 3), supporting the idea that Rad9 might affect BIR though a physical role at the DSB site.

**Fig. 2:**
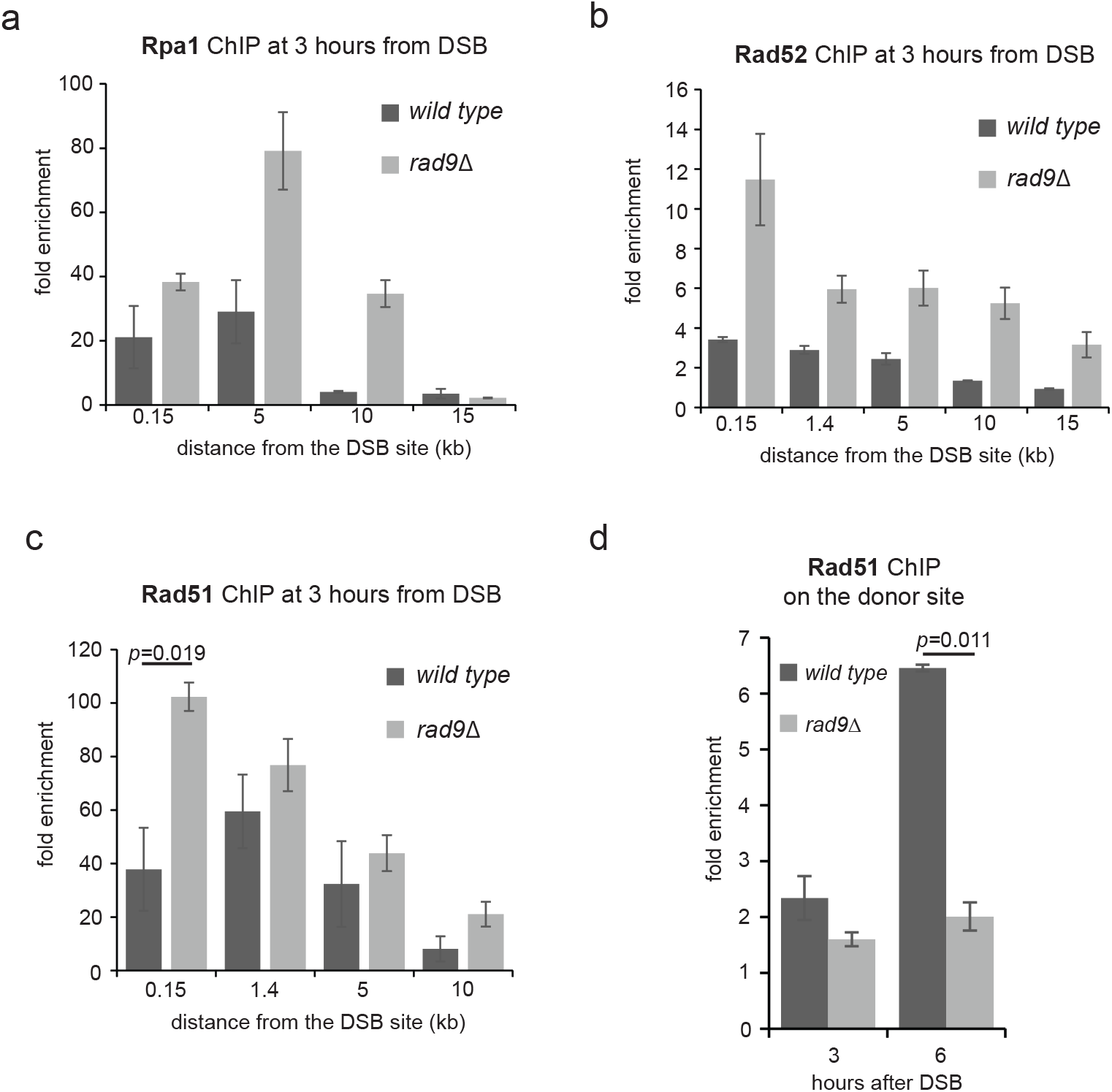
Rad9 limits Rpa1, Rad52 and Rad51 hyper-loading at the DSB site, and favours Rad51 binding at the donor site. ChIP analysis at the indicated positions near the cut site, after 3 hours since DSB induction, for the binding of Rpa1-Myc (a), Rad52-HA (b) and Rad51 (c), with SEM of three experiments. The indicated JKM strains were blocked in G2/M with nocodazole. The same experiment as in Extended data Fig. 2. d, Rad51 enrichment on donor site at Chromosome XI, 3h and 6h after DSB on Chromosome V in the indicated JRL92 strains, with SEM of three experiments. Cells were blocked in G2/M with nocodazole.

**Fig. 3:**
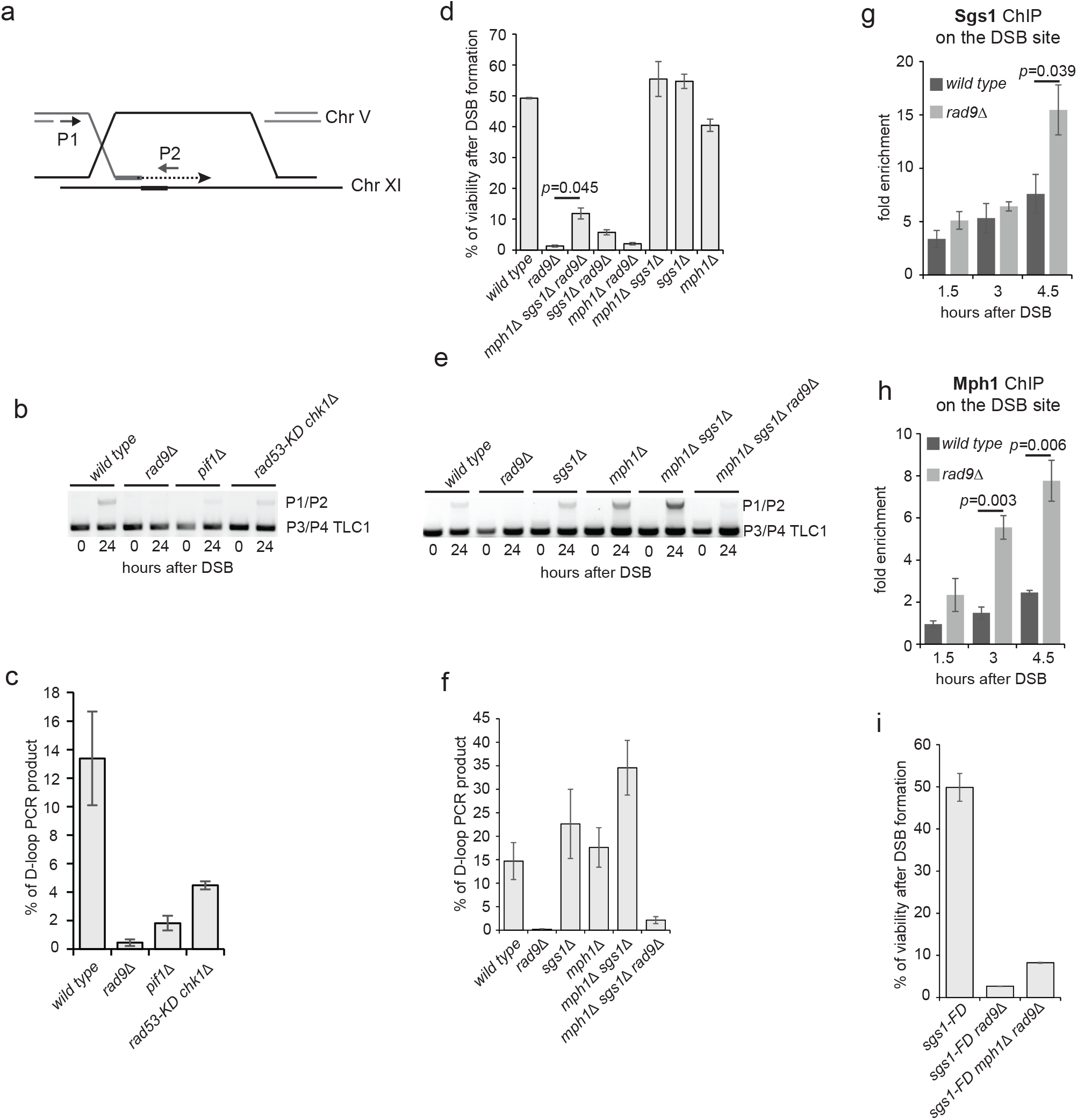
Rad9 promotes D-loop extension in BIR by limiting Sgs1 and Mph1. a, scheme of the PCR-based assay to test D-loop extension in JRL92 background. b, PCR-based primer extension assay to test D-loop in the indicated JRL92 strains. c, Quantification of the D-loop extension in three experiments as in (c), with SEM. d, BIR efficiency measured by cell viability in the indicated JRL92 strains, with SEM of three experiments. e, as in (b), with the indicated JRL92 strains. f, Quantification of the D-loop extension as in (e), with SEM of three experiments. g, Sgs1 binding at the DSB site in the indicated JKM strains blocked in G2/M with nocodazole, with SEM of three experiments. h, Mph1 binding at the DSB site in the indicated JKM strains blocked in G2/M with nocodazole, with SEM of three experiments. i, as in (d).

Despite the Rad51 binding at the DSB was increased (Fig. 2c), we found that it was not enriched at the donor site in *rad9*Δ, after DSB induction in G2/M blocked cells (Fig. 2d). This result indicates that cells lacking *RAD9* cannot form a stable Displacement-loop (D-loop) structure and synapsis between DSB and the donor template, providing molecular evidence of the failure in DSB repair by BIR. Consistent with these data, *RAD9* deletion impaired the D-loop extension, measured through a PCR-based assay to a greater extent than *rad*53-K227A *chk1*Δ and *pif1*Δ mutations (Fig. 3a, b, c). These results suggest that Rad9 promotes strand invasion and D-loop extension in BIR, through a mechanism independent on the Chk1 and Rad53 signalling. Of note, previous works have shown that the annealing between the two DNA strands in D-loop formation can either be promoted by a Rad51-dependent process or rejected by the Sgs1-Top3-Rmi1 complex and Mph1^11–15^. Strikingly, after deletion of the genes coding for the two helicases Sgs1 and Mph1 in *rad9*Δ cells, both viability and primer extension analyses showed a rescue to the level found in *pif1*Δ cells (Fig. 3d, e, f). Moreover, by ChIP analysis, we also found higher binding of Sgs1 and Mph1 on to a persistent DSB in *rad9*Δ cells (Fig. 3g, h). As Sgs1 interacts with Rpa1 and Rad51^16,17^, while Mph1 interacts with Rpa1 and Rad52^18,19^, we are tantalized to suggest that the hyper-loading of Sgs1 and Mph1 at the DSB may be related to the increased recruitment of Rad51, Rad52 and Rpa1 in *rad9*Δ cells (Fig. 2a, b, c, and Extended Fig. 1). In particular, it has been recently shown that Sgs1 interacts with Rad51 and the *sgs1*-F1192D (*sgs1*-FD) mutation abolishes this interaction^17^. We found that the *sgs1*-FD partially rescued the viability of cells lacking *RAD9* in the BIR assay (*rad9*Δ= 1.29 % ± 0.32; *sgs1*-FD *rad9*Δ= 2.67 % ± 0.03; *mph1*Δ *sgs1*-FD *rad9*Δ= 8.25 % ± 0.11) (Fig. 3d, i), similarly to the deletion of *SGS1* (*sgs1*Δ *rad9*Δ= 5.73 % ± 0.83; *mph1*Δ *sgs1*Δ *rad9*Δ= 11.87 % ± 1.78) (Fig. 3d). This result suggests that lowering the interaction between Sgs1 and Rad51 might reduce strand rejection, favouring BIR in *rad9*Δ cells. As because the interplay between Rad9 and Sgs1 regulates also DSB resection^5,6^, we measured the kinetic of ssDNA formation at different positions from the break in the *sgs1*-FD mutant. We found that this allele did not alter resection of an irreparable DSB, which still remained faster in the *sgs1*-FD *rad9*Δ double mutant cells (Extended data Fig. 2). These results, together with the viability of the *sgs1*-FD *rad9*Δ cells in the BIR assay (Fig. 3i), suggest that the kinetic of the DSB resection is not responsible per se of the severe BIR defect of the *rad9*Δ cells.

Because of the reduced levels of CO and long track gene conversion in *rad9*Δ cells (Fig 1a, b, c), we investigated the role of Rad9 in DSB repair by using an ectopic gene conversion (eGC) system^11^, which is coupled to DDC activation^20^. This assay allows us to detect both the CO and NCO repair products by Southern blotting (Fig 4a). Of note, deletion of *SGS1* and *MPH1* greatly increase CO^11,12^. Significantly, we found that *rad9*Δ cells had reduced repair product through both NCO and CO, with a major effect on the latter, when the DSB was induced in cells blocked in G2/M phase (Fig. 4b, c). Once again, by deleting *SGS1* and *MPH1* we observed a partial rescue of the DSB repair defect in *rad9*Δ cells, specifically for the less frequent class of repair associated with CO (Fig. 4b, c).

**Fig. 4:**
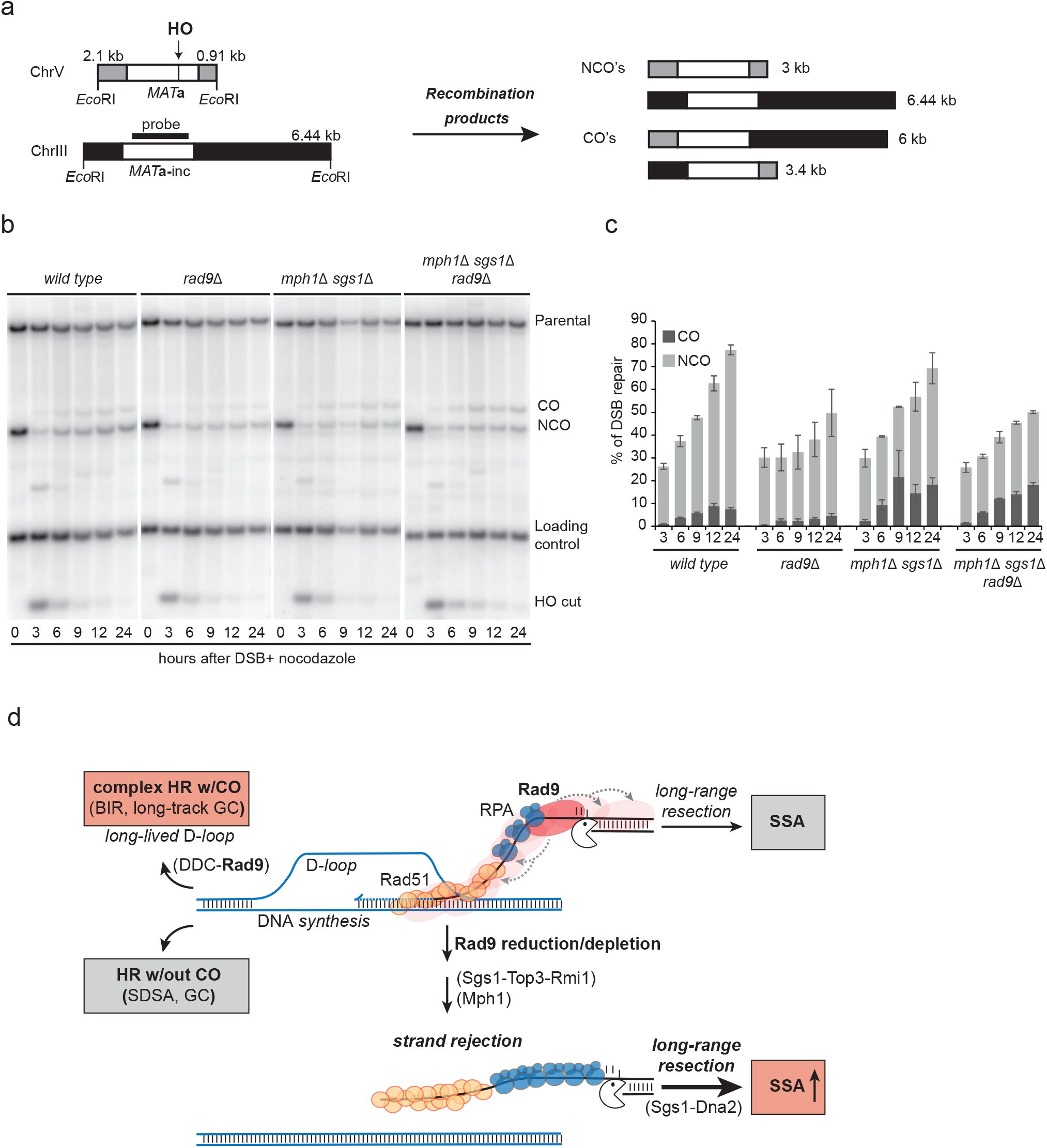
Rad9 promotes recombination events that require stable D-loop. a, scheme of the genetic system to test ectopic gene conversion (eGC) in tGI354 background. b, Southern blot of *Eco*RI-digested DNA to monitor DSB repair through eGC in the indicated tGI354 strains. CO, Crossover; NCO, Noncrossover. c, Densitometric analysis of the CO and NCO bands of three experiments as in (b), with SEM. d, working model: Rad9 controls HR, coordinating resection and strand rejection steps. After a DSB, Rad9 binding at the lesion limits: i) the Sgs1-Dna2 axis for the long-range resection and SSA; ii) the Sgs1-Top3-Rmi1 and Mph1 axes for the strand-rejection, favouring long-lived D loop and complex HR sub-pathways.

Overall, this work provides experimental evidence of an unprecedented role of Rad9 in DSB repair, controlling the fate of 3′ Rad51 filament in HR. Interestingly, limiting the Sgs1 helicase, Rad9 antagonizes the formation of extended 3′ ssDNA for recombination while, limiting both Sgs1 and Mph1, reduces the disassembling of D-loop structure once repair DNA synthesis is commenced (Fig. 4d). This reduces SSA and favours DSB repair through HR sub-pathways that require stable D-loop, such as long tract GC, eGC and BIR, also increasing the frequency of CO outcome. Accordingly, *RAD9* deletion limits sister chromatid exchanges and promotes Rad1/XPF-dependent translocations, likely through SSA ^21,22^. Moreover, in line with our model, eliminating the *SGS1* gene in *rad9*Δ cells causes dramatic levels of translocations between homeologous sequences and complex chromosomal rearrangements ^23,24^.

In conclusion, we suggest that the Rad9 recruitment at a DSB guarantees that the DNA break is engaged into complex and slow HR sub-pathways only when cell division is restrained by the DDC activation. This reduces the risk of premature segregation of chromosomes with DNA linkages caused by unresolved HR intermediates, which leads to the formation of anaphase bridges and deleterious chromosome rearrangements^25,26^. As such, this regulation might be critical to limit genetic instability, a hallmark of cancer cells. Indeed, 53BP1-depleted cancer cells accumulate ultrafine anaphase bridges, suggesting that this mechanism might be evolutionary conserved also in higher eukaryotes^27^.

## Methods

### Yeast strains, Media and Growth conditions

All the strains listed in Extended data Table 2 are derivative of JKM179, JRL92, tGI354 and W303. To construct strains standard genetic procedures were followed^28^. Deletions and tag fusions were generated by the one-step PCR system. The *sgs1-F1192D* was obtained using a Cas9 mediated gene targeting system^29^. For the indicated experiments, cells were grown in YEP medium enriched with 2% glucose (YEP+glu), 3% raffinose (YEP+raf) or 3% raffinose and 2% galactose (YEP+raf+gal). Unless specified all the experiments were performed at 28 °C. To block cells in G2/M, 20 μg/ml nocodazole was added to the cell culture.

### Cell viability assay

JRL92 derivative strains were inoculated in YEP+raf, grown O/N at 28°C. The following day, cells were normalized and plated on YEP+glu and YEP+gal. Plates were incubated at 28 °C for three days. Viability results were obtained from the ratio between number of colonies on YEP+gal and YEP+glu. Standard error of the mean (SEM) was calculated on two independent experiments.

### Southern blot analysis

Purified genomic DNA was digested with the appropriate restriction enzyme/s, probed with a specific ^32^P-labeled probe and scanned with a Typhoon Imager (GE healthcare). In JRL92 a *CAN1* fragment was used as a probe, the % of BIR repair has been calculated using the donor band as a loading control. Repair in tGI354 background was analyzed as described previously^20^. Densitometric quantification of the bands intensity was performed using the ImageJ software. The SEM was calculated on three independent experiments.

### ChIP analysis

ChIP analysis was performed as described previously^5^. Input and immunoprecipitated DNA were analysed by quantitative PCR, using a Bio-Rad CFX connect, or droplet digital PCR (ddPCR), using a Bio-Rad QX200 droplet reader. The oligonucleotides used are listed in Extended data Table 3. Data are presented as fold enrichment at the HO cut site (at the indicated distance from the DSB) over that at the *KCC4* locus on the left arm of chromosome III or *CAN1*locus on chromosome V. Then normalized to the corresponding input sample. SEM was calculated on two independent experiments.

### D-loop extension analysis

To measure the DNA synthesis after D-loop formation during BIR we adopted a strategy that was described previously^8^. In our experiments, the genomic DNA at 0h and 24h after DSB formation was amplified by PCR with oligonucleotides on the *CAN1* locus and, as a control, on *TLC1* locus (Expanded data Table 3). See a scheme in Fig. 2b.

### Quantitative analysis of DSB end resection by Real time PCR

Quantitative PCR (qPCR) analysis of DSB resection was performed as already described^30^. The oligonucleotides used are listed in Expanded data Table 2. The DNA was digested with the *Rsa*I restriction enzime (NEB) that cuts inside the amplicons at several distances from the DSB, but not in the *KCC4* control region on chromosome III. qPCR was performed on both digested and undigested templates using StoS Quantitative Master Mix 2X SYBR Green (Genespin) with the Bio-Rad CFX connect qPCR system. The ssDNA percentage over total DNA was calculated using the following formula: % ssDNA = *{100/ [(1 + 2^∆Ct^)/2]}/ f*, in which *∆Ct* values are the difference in average cycles between digested template and undigested template of a given time point and *f* is the HO cut efficiency measured by qPCR.

### DSB-induced recombination assay

Homozygous diploids of the wild type and *rad9*Δ strains derived from LSY2205-11C and LSY2543 (W303 background) were grown in YEP+raf overnight at 28°C. Cells were plated on YEP+glu and YEP+gal plates to calculate plating efficiency and ensure *ADE2* auxotrophy before I-*Sce*I induction. Separately, 2% galactose was added to the cell culture to induce I-*Sce*I. After 1.5h of induction, cells were plated on YEP+glu plates and incubated for 2-3 days at 28°C. Percentage of Red (the class of two long track conversions), White (the class of two short track conversions) and Red/White (the class of one long and one short track conversion) was determined only for recombinant colonies. All the plates were replicated on YEP+glu plates containing Hygromycin, Nourseothricin and Synthetic Complete (SC) medium-ura, SC-met, SC-ade, SC-ade+raf, SC-ade+gal plates to determine the percentage of recombinant colonies and distinguish the NCO, CO and BIR events in each sub-class of the colonies^7^.

### Statistical analysis

Statistical analysis was performed using Microsoft Excel or Prism softwares. p-values were determined by an unpaired two-tailed t-test. No statistical methods or criteria were used to estimate sample size or to include or exclude samples.

## Acknowledgements

We thank Lorraine S. Symington and James E. Haber for generously provided yeast strains. We thank Gerard Mazon for useful suggestions for using LSY2205-11C/LSY2543 background. We are grateful to James E. Haber and all the members of our laboratory for helpful discussions. This work was supported by grants to A.P. from Associazione Italiana Ricerca sul Cancro (AIRC_IG19917, AIRC_IG15488) and from Ministero Istruzione Università e Ricerca, MIUR (PRIN-2015LZE994). M.F. was supported by a fellowship from Fondazione Gabriella Dolfin Voyasidis-Accademia Nazionale dei Lincei. C.C.R. was supported by a fellowship from Università degli Studi di Milano (Assegno di ricerca –Tipo A).

## Author contributions

M.F. performed experiments in Fig. 1, 2, 3, 4 and Expanded data Fig. 1, 2, 3.

C.C.R. performed experiments in Fig 1b, c, and contributed to Fig. 2 and Fig. 3.

S.L. contributed to the ChIP analyses in Fig 2c and Fig. 3g, h, and Expanded data Fig. 1a.

A.P. supervised and coordinated all aspects of the work.

M.F., C.C.R. and A.P. wrote the manuscript.

---

**Extended data Fig. 1:**
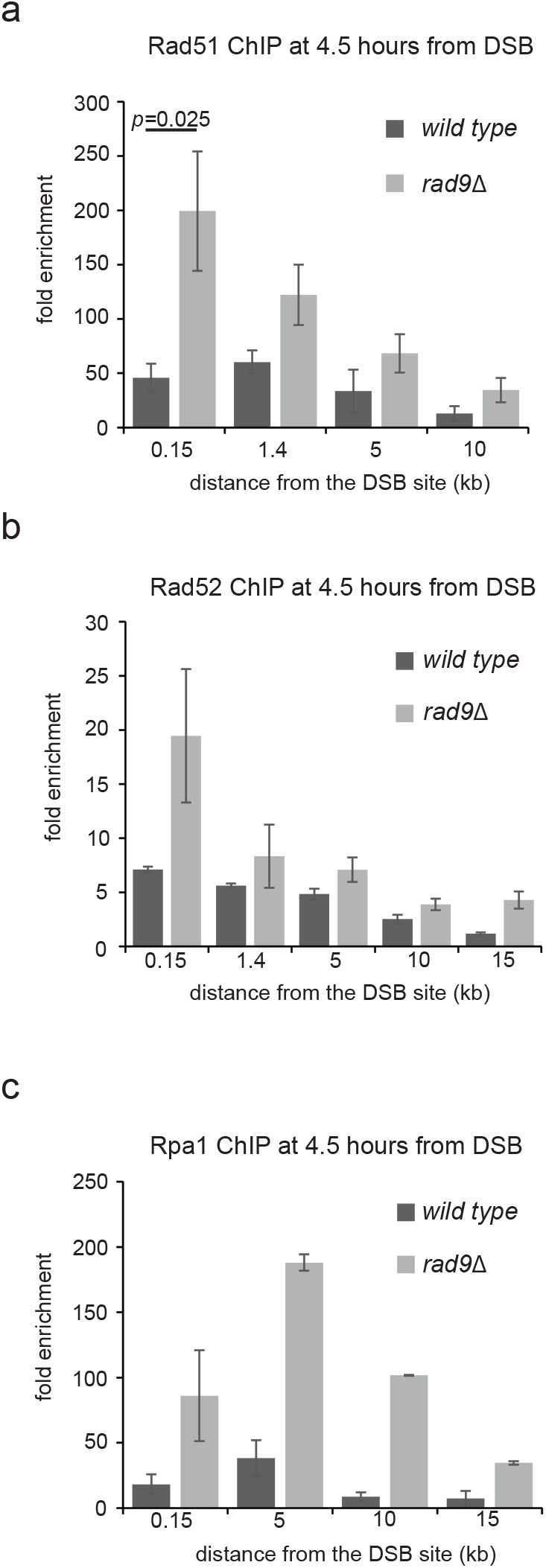
Rad9 limits Rpa1, Rad52 and Rad51 hyper-loading at the DSB site. ChIP analysis at the indicated positions near the cut site, 4.5 hours after DSB induction, for the binding of Rad51 (a), Rad52-HA (b) and Rpa1-Myc (c), with SEM of three experiments. The indicated JKM strains were blocked in G2/M with nocodazole. The same experiment as in Fig 2.

**Extended data Fig. 2:**
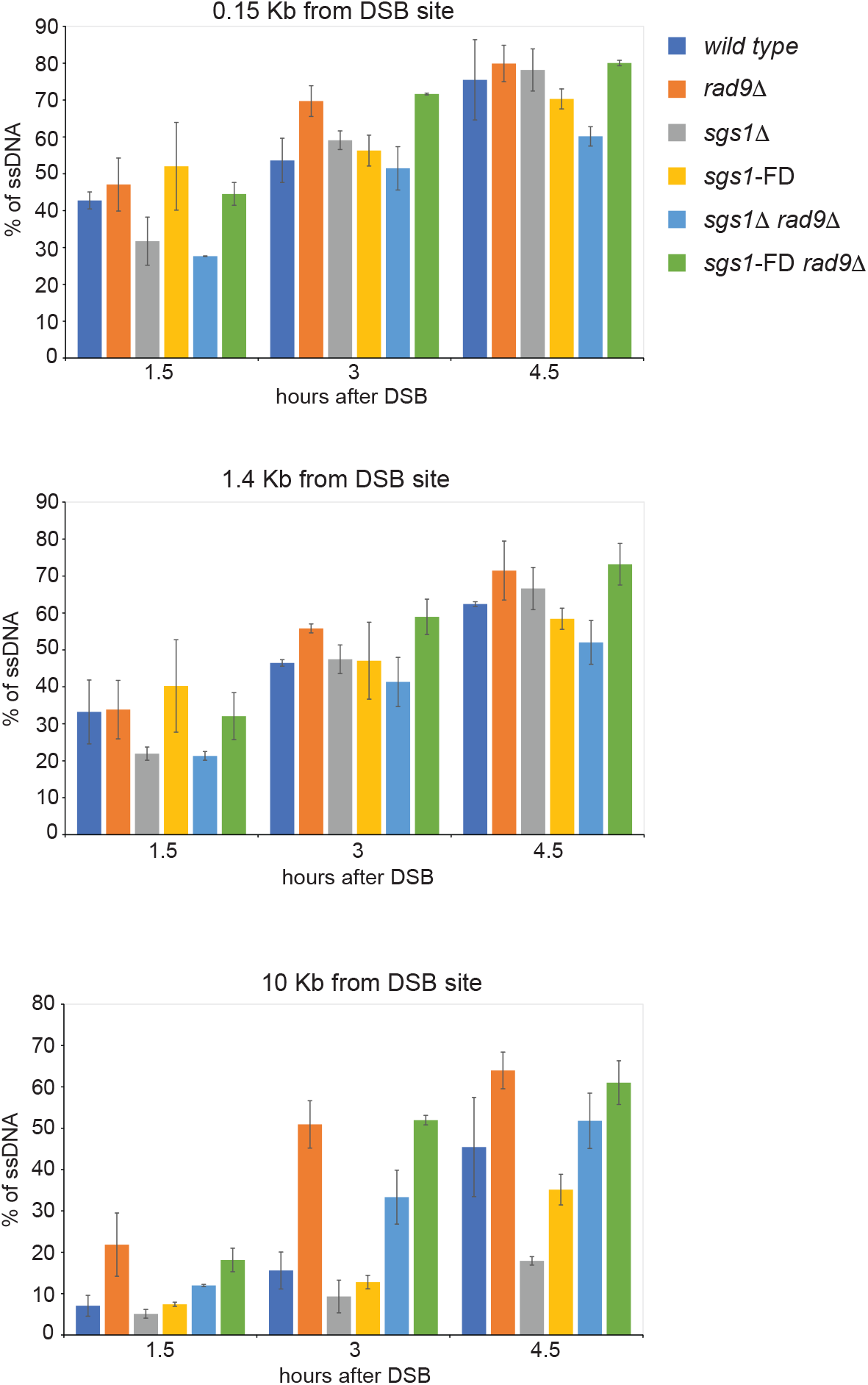
*rad9*Δ and *sgs1*-FD mutations do not alter DSB resection closed to the DSB. DSB resection analysis at 0.15 kb (a), 1.4 kb (b) and 10 Kb (c) from the DSB at the indicated times after HO induction, with SEM of three experiments. The indicated JKM strains were blocked in G2/M with nocodazole.

**Extended data Fig. 3:**
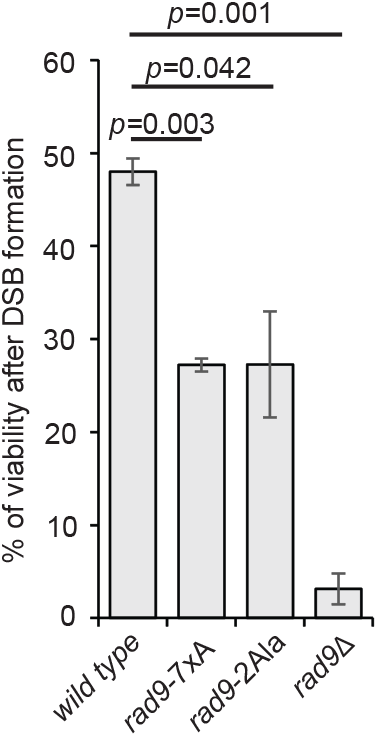
*rad9*-2Ala and *rad9*-7xA alleles partially rescue BIR of *rad9*Δ cells. BIR efficiency measured by cell viability in the indicated YRL92 strains, with SEM of two experiments.

**Extended data Table 1:** Distribution of colonies related to Fig. 1c.

|  |  | Diploid 1 | Diploid 1 | Diploid 2 | Diploid 2 |
| --- | --- | --- | --- | --- | --- |
|  |  | WT | *rad9Δ* | WT | *rad9Δ* |
| White/Red sectors | BIR | 4 | 3 | 5 | 4 |
|  | BIR-like | 1 | 0 | 4 | 0 |
|  | CO | 6 | 2 | 6 | 4 |
|  | Unclassified LOH | 0 | 0 | 0 | 0 |
|  | NCO | 40 | 18 | 68 | 13 |
| Red colonies (Recombinants) | BIR | 0 | 0 | 3 | 1 |
|  | BIR-like | 5 | 0 | 0 | 0 |
|  | CO | 7 | 2 | 6 | 2 |
|  | Unclassified LOH | 1 | 13 | 3 | 8 |
|  | NCO | 105 | 35 | 64 | 31 |
| White colonies | BIR | 0 | 0 | 0 | 0 |
|  | BIR-like | 0 | 1 | 0 | 0 |
|  | CO | 3 | 1 | 5 | 0 |
|  | Unclassified LOH | 13 | 8 | 10 | 11 |
|  | NCO | 91 | 171 | 72 | 150 |
| Red (Non recombinants) |  | 12 | 5 | 7 | 5 |
| Total colonies |  | 288 | 259 | 253 | 229 |
| Total recombinants |  | 276 | 254 | 246 | 224 |

**Extended data Table 2:**
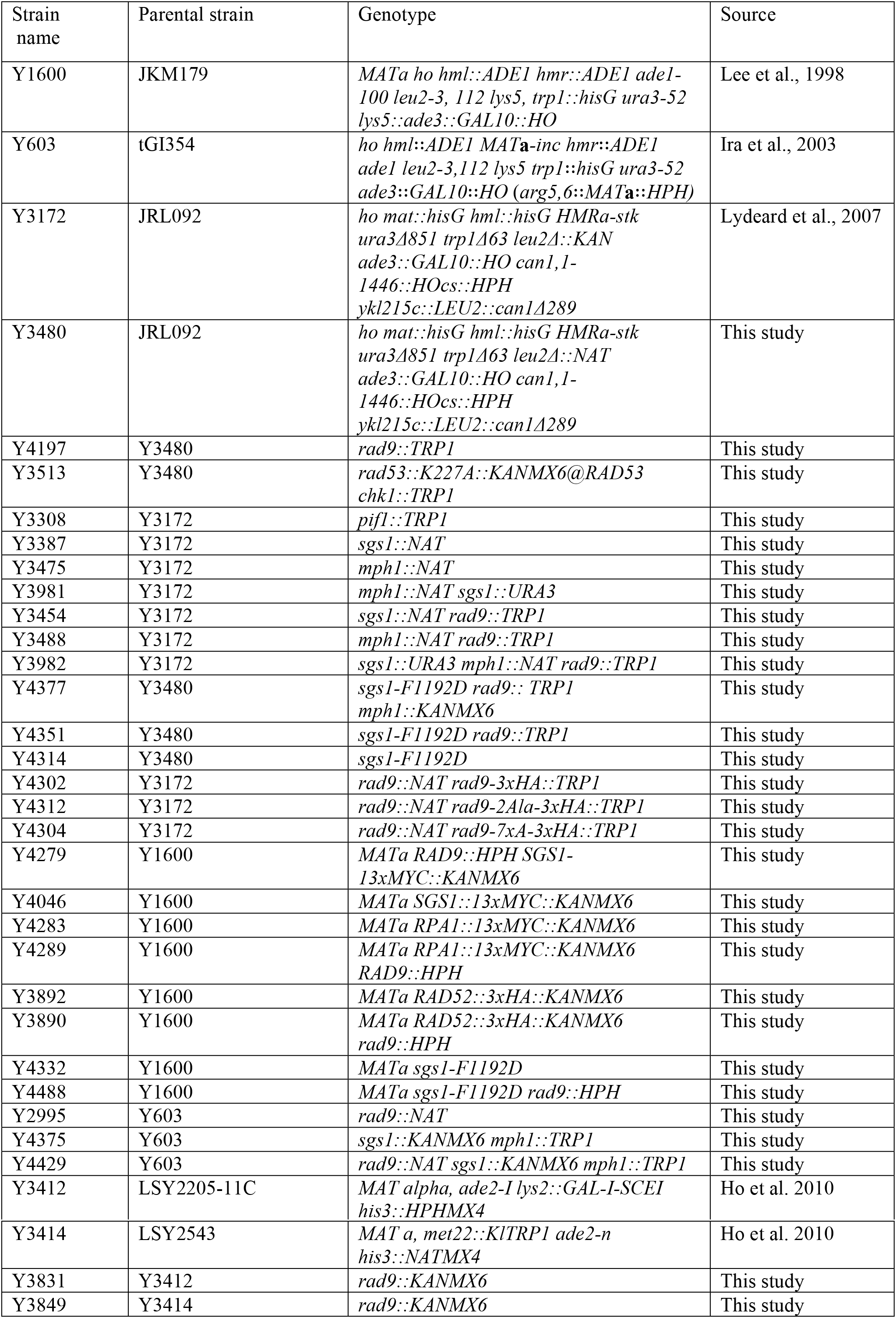
List of the yeast strains used in this study.

**Extended data Table 3:**
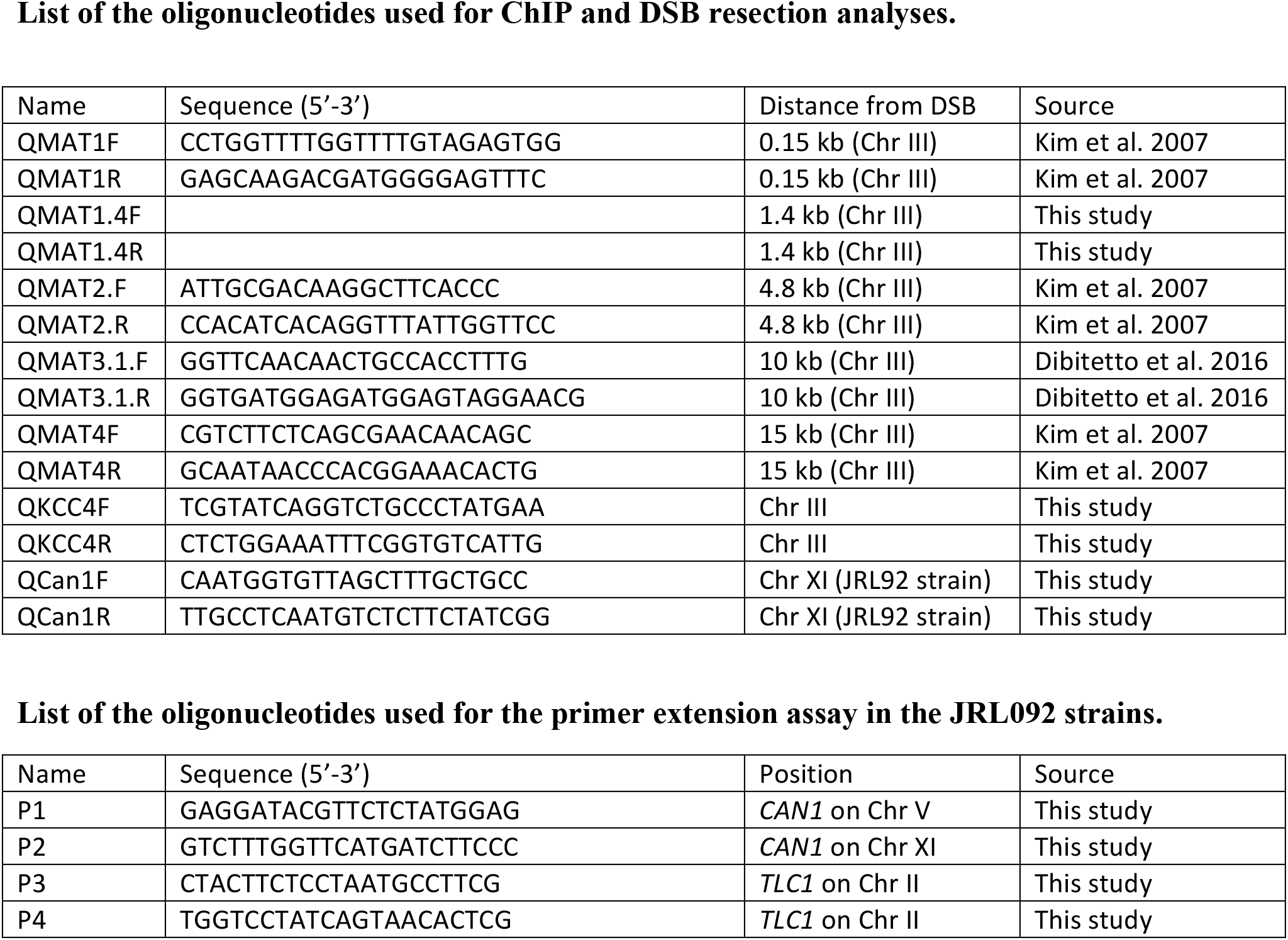
List of the oligonucleotides used in this study.

